# The human experience with intravenous levodopa

**DOI:** 10.1101/024794

**Authors:** Shan H. Siddiqi, Natalia K. Abraham, Christopher L. Geiger, Morvarid Karimi, Joel S. Perlmutter, Kevin J. Black

## Abstract

**Objective:** To compile a comprehensive summary of published human experience with levodopa given intravenously, with a focus on information required by regulatory agencies.

**Background:** While safe intravenous use of levodopa has been documented for over 50 years, regulatory supervision for pharmaceuticals given by a route other than that approved by the U.S. Food and Drug Administration (FDA) has become increasingly cautious. If delivering a drug by an alternate route raises the risk of adverse events, an investigational new drug (IND) application is required, including a comprehensive review of toxicity data.

**Methods:** Over 200 articles referring to intravenous levodopa (IVLD) were examined for details of administration, pharmacokinetics, benefit and side effects.

**Results:** We identified 142 original reports describing IVLD use in humans, beginning with psychiatric research in 1959-1960 before the development of peripheral decarboxylase inhibitors. At least 2781 subjects have received IVLD, and reported outcomes include parkinsonian signs, sleep variables, hormones, hemodynamics, CSF amino acid composition, regional cerebral blood flow, cognition, perception and complex behavior. Mean pharmacokinetic variables were summarized for 49 healthy subjects and 190 with Parkinson’s disease. Side effects were those expected from clinical experience with oral levodopa and dopamine agonists. No articles reported deaths or induction of psychosis.

**Conclusion:** At least 2781 patients have received i.v. levodopa with a safety profile comparable to that seen with oral administration.

## Introduction

Parkinson disease (PD), the second most common neurodegenerative disease, is associated with impairments in dopaminergic neurotransmission in the basal ganglia. Replacement of dopamine has been the cornerstone of treatment for PD. Because dopamine itself does not cross the blood-brain barrier (BBB), its immediate precursor levodopa (L-3,4-dihydroxphenylalanine, L-DOPA) is adminstered since it does cross the BBB (Hornykiewicz, 1963; Cotzias et al., 1967; Birkmayer and Hornykiewicz, 2001). Although purified levodopa was first ingested by mouth in 1913 (Roe, 1997), it was first used for medical treatment by intravenous rather than oral administration (Pare and Sandler, 1959; Birkmayer and Hornykiewicz, 2001).

Oral levodopa has become the preferred method of treatment clinically, but intravenous levodopa administration still holds advantages over the oral form for some research studies. First, the rapid administration of intravenous levodopa is often necessary for certain study designs, including those focused on the pharmacokinetics and pharmacodynamics of the drug. Additionally, intravenous administration leads to more predictable plasma levodopa concentration because oral medications have highly variable absorption characteristics, especially in PD patients (Bushmann et al., 1989), with differences in absorption based on sex and age (Robertson et al., 1989; Kompoliti et al., 2002). Intravenous levodopa permits researchers to keep brain levodopa concentrations constant while assessing physiological responses over time. Furthermore, intravenous levodopa has sometimes been used clinically in patients who cannot tolerate oral medications, such as PD patients during surgery or on total parenteral nutrition.

Current U.S. FDA regulations focus heightened scrutiny on research in which drugs are delivered by a route for which the drug has not been approved. Predictably, in addition to any safety benefits, the heightened scrutiny has also created practical obstacles to research with intravenous levodopa, as described for instance by Rascol and colleagues (2001, p. 250). Specifically, an IND (Investigational New Drug) application must be submitted if risks of intravenous levodopa are significantly higher than those of oral levodopa (<u>§21 CFR 312.2(b)(iii)</u>). Therefore, the overall goal of this paper is to facilitate research use of IV levodopa by compiling a literature review that comprehensively summarizes the human experience with intravenously administered levodopa. We tabulate the extent of human exposure, side effects, benefits, and efficacy. We also summarize pharmacokinetic (PK) and pharmacodynamic (PD) parameters from these studies.

## Methods

The authors searched MEDLINE and OVID, reviewed selected books, searched toxicity databases, and followed references cited in those sources. Articles written completely in languages other than English, German, Italian, Spanish, or Portuguese were excluded. Search terms included (levodopa / L-dopa / DOPA) AND (intravenous / intravascular/ infusion / injection / i.v.); limit to humans; search date through May, 2015. Studies using oral or intraduodenal L-DOPA administration were excluded except for PK/PD studies cited in Table 2. Studies in which IV levodopa was always coadministered with monoamine oxidase inhibitors (MAOIs) or catechol-O-methyltransferase (COMT) inhibitors were excluded. Levodopa methyl ester (Juncos et al., 1987) and D,L-dopa (Pare and Sandler, 1959) were included, but PK/PD calculations were corrected for the difference in molecular weights. Co-administered drugs were reported if included by the authors.

We recorded total dose and maximum infusion rate. We also recorded pharmacokinetic (PK) and pharmacodynamic (PD) parameters where available, including steady state volume of distribution (VOD), clearance, distribution half life (*t*_½α_), elimination half life (*t*_½_ or *t*_½β_), *E*_*max*_, and *EC*_50_. Reported data were used to calculate any missing PK parameters where possible. Additionally, any reports on efficacy were noted. Side effect frequency was recorded if reported. The number of subjects and subject conditions (Parkinson disease, other disease states or healthy volunteers) were recorded for each study. Average PK parameters were calculated across studies, weighted by the number of subjects.

## Results

142 articles reporting intravenous levodopa administration were identified. Most subjects with parkinsonism were diagnosed with idiopathic PD, but some studies reported a variety of etiologies including postencephalitic and vascular parkinsonism and PSP. PD patients differed in their history of prior drug treatment before the studies with conditions including de novo, fluctuating, on-off, and stable. Some subjects were treated with levodopa for conditions other than PD (see Table 1: Patient Populations and Response Parameters), including other movement disorders (dystonia, progressive supranuclear palsy [PSP], neuroleptic malignant syndrome [NMS], primary psychiatric disorders (schizophrenia, mood disorders, personality disorders), endocrine disorders (diabetes mellitus, essential obesity, hypopituitarism), hepatic disease (alcoholic cirrhosis, steatohepatitis, hepatic encephalopathy), cardiac valvular disease, and asthma. Healthy controls were also included in some studies.

**Table 1:** Patient populations and response parameters. Subject populations given intravenous levodopa and responses to drug measured in studies listed in Table 3.

| Patient populations | Response Parameters |
| --- | --- |
| Healthy volunteers | Vital signs: |
| Movement Disorders: | Heart rate, blood pressure, temperature, respirations |
| Parkinson's (de novo, stable, fluctuators, on-off) | Cardiovascular: |
| Progressive supranuclear palsy | ECG |
| Parkinson's disease psychosis | Cerebral blood flow |
| Carcinoma of the Rectum | Renal: |
| Stereotactic surgery | Urine flow |
| Post-menopausal women | Urinary sodium excretion |
| Tourette Syndrome/tic disorders | Potassium excretion |
| Asthma | Plasma renin activity |
| Schizophrenia | Renal plasma flow |
| Mood disorders: | Metabolism: |
| Mild to moderate depression | Urinary metabolite excretion |
| Treatment-resistant depression | Cerebral metabolism |
| Bipolar depression | Plasma metabolites |
| Cyclothymic disorder | CSF amino-acid composition |
| Borderline personality disorder | PD motor improvement |
| Neuroleptic Malignant Syndrome | UPDRS, walking, tapping, etc. |
| Hepatic disorders: | Dyskinesias |
| Alcoholic cirrhosis | Tic improvement |
| Steatohepatitis | Neuropsychiatric:: |
| Hepatic Encephalopathy | Cognition |
| Endocrine Disorders: | Mood |
| Diabetes mellitus | Behavior |
| Essential obesity | Psychosis |
| Hypopituitarism | Dementia |
| Cardiovascular Disease: | EEG (including REM sleep EEG) |
| Atrial septal defect | Endocrine: |
| Rheumatic Valvular disease | Prolactin, HGH, ACTH, LH, Vasopressin |

Pharmacokinetic data were reported for a total of 251 human subjects (see Table 2: Pharmacokinetics of Levodopa). Co-administration of a peripheral decarboxylase inhibitor (PDI) lowered the clearance and elimination half-life of intravenously administered levodopa, while there was no notable effect of PDIs on volume of distribution. Additional PK data are available from studies that gave levodopa by other routes (Sasahara et al., 1980a; Poewe, 1993; Muhlack et al., 2004; LeWitt et al., 2009), and several studies report on the bioavailability of oral doses relative to intravenous administration (Sasahara et al., 1980b; Robertson et al., 1989; Kompoliti et al., 2002).

**Table 2:** Pharmacokinetics of levodopa. Summary of pharmacokinetic parameters with weighted means.

| Reference | Patient Group | Clearance |  | Volume of distribution |  | Elimination half-life |  | Distribution half-life |  |
| --- | --- | --- | --- | --- | --- | --- | --- | --- | --- |
|  |  | n | Mean (L/kg/hr) | n | Mean (L/kg) | n | Mean (hr) | n | Mean (hr) |
| (Birkmayer et al., 1973) | PD | 50 | 1.61 | 50 | 2.44 | 50 | 1.05 |  |  |
| (Bredberg et al., 1990) | Fluctuating | 5 | 0.37 |  |  |  |  |  |  |
| (Chan et al., 2004) | De Novo | 12 | 0.36 | 12 | 0.63 | 12 | 2.25 | 12 | 0.17 |
|  | Chronic | 12 | 0.35 | 12 | 0.49 | 12 | 1.47 | 12 | 0.17 |
| (Durso et al., 2000) | "Slow" CD Absorption |  |  |  |  | 5 | 1.18 |  |  |
|  | "Rapid" CD Absorption |  |  |  |  | 4 | 1.15 |  |  |
| (Fabbrini et al., 1987) | De Novo | 4 | 0.13 | 4 | 0.26 | 4 | 1.44 |  |  |
|  | Stable | 6 | 0.11 | 6 | 0.22 | 6 | 1.41 |  |  |
|  | Wearing-Off | 6 | 0.13 | 6 | 0.30 | 6 | 1.67 |  |  |
|  | On-Off | 12 | 0.13 | 12 | 0.30 | 12 | 1.54 |  |  |
| (Hardie et al., 1986)<sup>a</sup> | Fluctuating | 7 | 1.14 | 7 | 2.63 | 7 | 1.60 | 7 | 0.13 |
| (Gancher et al., 1987)<sup>b</sup> | De Novo | 5 | 0.34 | 5 | 0.56 | 5 | 1.70 | 5 | 0.10 |
|  | Stable | 4 | 0.33 | 4 | 0.62 | 4 | 1.80 | 4 | 0.11 |
|  | Fluctuating | 11 | 0.32 | 11 | 0.65 | 11 | 2.00 | 11 | 0.10 |
| (Nutt et al., 1985) | 2 hr IV | 7 | 0.55 | 7 | 0.67 | 7 | 1.38 | 7 | 0.07 |
| (all PD, fluctuating) | 2 hr IV + PDI | 7 | 0.30 | 7 | 0.80 | 7 | 2.01 | 7 | 0.11 |
|  | ≥20 hr IV | 4 | 0.52 | 4 | 0.88 | 4 | 1.19 | 4 | 0.11 |
|  | ≥20 hr IV + PDI | 4 | 0.28 | 4 | 1.09 | 4 | 2.60 | 4 | 0.33 |
| (Nutt et al., 1992) | De Novo | 8 | 0.44 | 8 | 0.75 | 8 | 1.60 |  |  |
|  | Stable | 12 | 0.42 | 12 | 0.75 | 12 | 1.70 |  |  |
|  | Fluctuating | 9 | 0.39 | 9 | 0.63 | 9 | 1.50 |  |  |
| (Poewe, 1993)<sup>e</sup> | PD |  | 1.40 |  | 3.00 |  | 1.50 |  | 0.09 |
| (Roberts et al., 1995)<sup>c,d</sup> | Healthy | 8 | 0.37 | 8 | 1.13 | 8 | 2.15 |  |  |
|  | Healthy + selegine | 8 | 0.37 | 8 | 2.01 | 8 | 3.78 |  |  |
| (Robertson et al., 1989)<sup>c</sup> | Healthy elderly | 9 | 0.85 | 9 | 1.01 | 9 | 0.82 |  |  |
|  | Healthy young | 8 | 1.40 | 8 | 1.65 | 8 | 0.82 |  |  |
|  | Healthy elderly + PDI | 8 | 0.35 | 8 | 0.62 | 8 | 1.23 |  |  |
|  | Healthy young + PDI | 8 | 0.56 | 8 | 0.93 | 8 | 1.16 |  |  |
| (Sasahara et al., 1980b) | PD | 5 | 1.38 | 5 | 1.29 | 5 | 0.65 |  |  |
| (Stocchi et al., 1992) | Intravenous Bolus | 6 | 0.97 | 6 | 0.96 | 6 | 0.83 |  |  |
| (all "on-off") | Intravenous Infusions | 2 | 0.63 | 2 | 0.82 | 2 | 0.90 |  |  |
|  | **Total n** | 212 |  | 242 |  | 251 |  | 73 |  |
|  | **Weighted mean** |  | **0.719 L/kg/hr** |  | **1.18 L/kg** |  | **1.50 hr** |  | **0.14 hr** |

**Notes:**
a assumed mean weight to be 70 kg for VOD
b values read from graphs
c half-life estimated from relationship: clearance = (ln 2 \* VOD)/ elim. T1/2
d assumed mean weight to be 70 kg for CL
e unpublished data, not included in weighted mean calculations of pharmacokinetic parameters

The pharmacodynamic data (see Table 3: Reports of Human Experience with IV Levodopa) obtained from the literature surveyed a total of 2651 human subjects, with a significant variety of patient groups studied and a multitude of response parameters (see Table 1). From these articles, no side effects were reported for a total of 1260 subjects. The highest total dose was 4320 mg in one day, given to a patient with idiopathic PD and carcinoma of the retina. The patient reported no side effects or adverse effects at this dose. The highest single bolus dose was 200 mg, while the highest infusion rates were 5.0 mg/kg/hr.

**Table 3:** The human experience with intravenous levodopa. Summary of published studies reporting intravenous levodopa use in humans, 1959 to early 2015.

| Reference | N | Diagnosis | PDI | Concomitant Drugs | Total Dose | Maximum Rate | Side Effects/Comments |
| --- | --- | --- | --- | --- | --- | --- | --- |
| (Abramsky and Goldschmidt, 1974) | 4 | acute hepatic encephalopathy in cirrhotic patients with gastrointestinal bleeding | none mentioned | none mentioned | for several days (between 3 to 5 days depending on the patient) | 600-1200 mg/day | Levodopa was administered intravenously with striking and rapid improvement of the comatose state. Within 2-5 hours the patients had recovered their normal mental state. |
| (Aebert, 1967) | 11 | 10 PD, 1 post-encephalitis lethargica | none mentioned | none mentioned | 75-1375 mg | 75-100 mg/10-15 min | No side effects mentioned. |
| (Argyelan et al., 2008) | 15 | PD | none mentioned | none mentioned | not given | 0.83 mg/kg/hr | No side effects mentioned. Levodopa was associated with increases in learning-related activation in the left dorsal premotor cortex and in the right pre-supplementary motor area. In the former region, there was recovery of the normal activation response by levodopa. In the latter region, there was a treatment-mediated gain of response in that significant learning-related activation was present only when the patients were scanned on levodopa therapy. |
| (Baldy-Moulinier et al., 1977) | 19 | 12 alcoholic hepatic cirrhosis and hepatic encephalopathy; 3 alcoholic hepatic cirrhosis; 3 fatty liver (alcoholic) without cirrhosis; 1 healthy | none mentioned | none mentioned | 125 mg | 125 mg bolus | No effects on EEG, ECG, humeral arterial pressure, rectal temperature, cerebral perfusion or metabolism at this dose. |
| (Bara-Jimenez et al., 2003) | 15 | moderate to advanced PD | carbidopa | KW-6002 (Adenosine A2A receptor antagonist) | infusion of "optimal dose levodopa" | 725±65 ng/mL | No side effects mentioned for L-dopa plus placebo. There were no drug-related serious adverse events. Levodopa plus KW-6002 appeared generally safe and well-tolerated. |
| (Baronti et al., 1992) | 9 | moderate to severe PD (III to V) | carbidopa | Terguride (dopamine agonist); domperidone in 4 subjects | Variable, 26-55 mg/hr (from 5:00 AM until end of day's study) | 55 mg/hr | No side effects noted for L-dopa alone. For terguride plus levodopa, subjects had mild, transient asymptomatic orthostatic hypotension, headache, nausea, nervousness, drowsiness, light-headedness, and epigastric distress. |
| (Birkmayer and Hornykiewicz, 1962) | not given | not given | none mentioned | none mentioned | 50-150 mg | 150 mg | No side effects mentioned. |
| (Birkmayer and Hornykiewicz, 1964) | 200 | not given | none mentioned | none mentioned | 25 mg, once or twice a week, for up to 3 years | "slow infusion" | Unclear whether L-dopa was administered without MAO inhibitors or nialamide. Nausea, vomiting and fainting were the major side effects which reversely correlated with the level of benefit. |
| (Birkmayer and Hornykiewicz, 1962) | 132 | PD | none | MAO inhibitor (Ro-4/2637), caffeine, or ephyllin. | 50-150 mg infusions twice a week for two weeks. | 150 mg | L-dopa caused nausea and vomiting, if combined with MAO inhibitor. Caffeine or Ephyllin could reduce L-dopa side effects. |
| (Birkmayer and Mentasti, 1967) | 15 | PD | Ro 4-4602 (benserazide) | none mentioned | 50 mg | 50 mg | No side effects mentioned. Decarboxylase inhibitor increased the benefit of L-dopa. |
| (Birkmayer, 1967) | 1 | PD | none mentioned | none mentioned | 50 mg | not mentioned | No side effects mentioned. |
| (Birkmayer and Hornykiewicz, 1961) | 20 | parkinsonism (PD, postencephalitic parkinsonism, and vascular parkinsonism) | none mentioned | none mentioned | up to 150 mg | "slow i.v." (cite Degkwitz et al.) | No side effects mentioned. |
| (Black et al., 2003) | 127 | 55 PD, 20 chronic tic disorders, 52 normal | carbidopa | none mentioned | 2.2 mg/kg | 1.735 mg/kg /10 min | In healthy patients at high doses: nausea, vomiting, feeling uncomfortably hot, increased pulse rate. In PD patients at high doses: no side effects. In healthy patients at intermediate doses: nausea, vomiting. In PD patients at intermediate doses, some had dyskinesias but no nausea or vomiting. At low doses: there was some nausea in healthy patients. |
| (Black et al., 2010b; Black et al., 2010a) | 21 | PD | carbidopa | tozadenant (SYN115) | 0.6426 mg/kg | $2.882 \times 10^{-5} \times (140 - \text{age})$ mg/kg/min | Carbidopa 200mg was given by mouth at least an hour before the levodopa infusion began, using the method of (Gordon et al., 2007) and a target plasma concentration of 600ng/ml. |
| (Blanchet et al., 1999) | 8 | PD (postmenopausal women with mild to moderate PD) | carbidopa | Estradiol | 29± 4mg/10min twice per day | 33 mg/10 min | The threshold dose of levodopa necessary to provide definite antiparkinsonian efficacy was reduced significantly by 17[beta]-estradiol from 29 to 21 mg. |
| (Braun et al., 1987) | 7 | idiopathic PD | carbidopa | SKF38393 (selective D-1 agonist) administered orally in double blind, placebo-controlled, crossover design. | (10 to 80 mg/hr) x 12 hr | 80 mg/hr | No dyskinesias occurred with levodopa and simultaneous SKF 38393 treatment. Dyskinesias at higher, supraoptimal doses. No side effects mentioned for L-Dopa alone: no orthostatic changes in blood pressure; patients remained asymptomatic throughout. Hematological parameters and blood chemistries remained within normal limits. |
| (Bredberg et al., 1990) | 5 | PD (advanced) | benserazide | none mentioned | not given | 1.5mg/min | No side effects mentioned. |
| (Brod et al., 2012) | 12 | PD | carbidopa | none | 2 mg/kg | 1 mg/kg/hr | Study compared low doses of carbidopa to higher doses. Side effects mostly related to parkinsonian symptoms associated with lower dose of IV levodopa than the patient's usual oral dose. |
| (Bronaugh et al., 1975) | 21 | PD (15 idiopathic, 2 secondary to encephalitis lethargic, 2 associated with progressive supranuclear palsy) | none mentioned | none mentioned | calculated:30.8 - 56µg (for 7 patients, and for 6 patients who were already on 3.0g/day orally) | 7.7-14µg/4hr on top of an oral dose of 3.0g/day | No side effects mentioned. Percent conjugation of L-dopa and metabolites given. |
| (Bruck et al., 1965) | 20 | 10 PD, 10 healthy | none mentioned | none mentioned | 100 mg for PD, 50 mg for healthy individuals | 50-100 mg/20-30 min | Nausea, lightheadedness, syncope, unpleasant sensation in head and abdomen, and increased BP by 10-20 mmHG. |
| (Bruno and Brigida, 1965) | 18 | Schizophrenia | none mentioned | Haloperidol | 100-170 mg | 2 mg/kg/5min | No side effects mentioned for L-Dopa alone, only in combination with Haloperidol. |
| (Bruno and Bruno, 1966) | 40 | Schizophrenia | none mentioned | 20 received haloperidol, 20 received chlorpromazine | 2 mg/kg | 2 mg/kg/5 min | Neuroleptic-induced parkinsonism improved in both groups. Some improvement in antipsychotic-induced negative symptoms. Some patients developed nausea/vomiting, sweating, warmth/flushing, and dizziness (quantity not reported). No significant change in pulse or blood pressure. |
| (Camicioli et al., 2001) | 5 | PD (idiopathic), functionally independent | carbidopa | Methylphenidate (in one trial, compared to levodopa alone) | 2 mg/kg | 2 mg/kg/hr | Apart from bothersome dyskinesias in one patient, patients did not report side effects or difficulties with treatments. No effect on HR or diastolic blood pressure, but L-Dopa led to a drop in systolic blood pressure. The drop in systolic blood pressure caused by L-Dopa was reduced when methylphenidate was administered alongside L-Dopa. No changes in mood, anxiety, arousal, or concentration before or after medications. Motor UPDRS scores were improved, as were tapping rates for both sides and both walking steps and rate. |
| (Chan et al., 2004) | 25 | idiopathic PD | carbidopa | none mentioned | 2.35 g/day (x 3 days) | 1.6mg/kg/hr (x 2hrs x 3 days) | No side effects mentioned. |
| (Chung et al., 2005) | 14 | idiopathic PD | carbidopa | Paroxetine | 2.0 mg/kg/day x 4 weeks | 1.0 mg/kg/hr | No side effects mentioned. No serious adverse effects. |
| (Chung et al., 2010) | 22 | PD (15 with levodopa-induced dyskinesia) | carbidopa | none mentioned | 2-3 mg/kg | 1.5 mg/kg/hr | No side effects mentioned. |
| (Davis et al., 1991) | 10 | idiopathic PD | carbidopa | none mentioned | none mentioned: just found "optimal dose rate" Total of 4 consecutive doses at the optimal rate were given, so highest total dose was 4.4 mg/day | 1.1 mg/kg/10 min | Modest worsening of motor scores after levodopa stopped. Patients with unpredictable motor fluctuations have higher requirements for levodopa, both orally and intravenously, compared to those with simple wearing-off phenomena. |
| (Degkwitz et al., 1960) | ≥ 22 | Psychiatric patients and normal controls | none | none mentioned | 50-350 mg | bolus (at least, ≤10 min) | No side effects mentioned. |
| (Durso et al., 1997) | 8 | idiopathic PD | carbidopa | none mentioned | 150 mg bolus | 150 mg bolus stable isotope-labeled LD/12-15 minutes | No side effects mentioned. |
| (Durso et al., 2000) | 9 | idiopathic PD | carbidopa | none mentioned | 150 mg bolus | 150 mg bolus <sup>13</sup>C labelled L-DOPA/12-15 minutes | Average reduction in systolic blood pressure was 22 mmHg (14, 10-40). No prolonged cardiac arrhythmias were noted during infusion or subsequent 6-hour monitoring. |
| (Fabbrini et al., 1987) | 28 | idiopathic PD | carbidopa | none mentioned | 1.5 mg/kg/hr for ≥ 16 hr | 1.5mg/kg/hr | No side effects mentioned. |
| (Fabbrini et al., 1988) | 48 | idiopathic PD | carbidopa | none mentioned | 19.2 mg/kg | 2.0 mg/kg/hr | No side effects mentioned. |
| (Fasano et al., 1970a) | 66 | PD | benserazide 150mg i.v. | stimulant ("amfetamino-simile") | not stated | not stated | The authors say, "no side effects were reported" with i.v. levodopa, whereas chronic oral levodopa dosing (without benserazide) produced side effects in 87% of patients ("psychic disturbances," dyskinesias, nausea, vomiting, and orthostatic hypotension). |
| (Fasano et al., 1970b) | 75 | PD | none mentioned | none mentioned | not stated | not stated | No side effects mentioned. |
| (Fehling, 1966) | 25 | PD | none | none mentioned | 1.5 mg/kg | 1.5 mg/kg over 13 minutes (6.9 mg/kg/hr) | Levodopa did not differ from placebo in terms of clinical improvement. Levodopa caused a brief period of nausea in 9 patients and vomiting in 2 patients. Levodopa and placebo did not differ in their effects on blood pressure. |
| (Feigin et al., 2001) | 7 | PD | none mentioned | none mentioned | not given | 100 mg/hr (mean, 67.1 ± 25.6 mg/hr) | No side effects mentioned. |
| (Feigin et al., 2002) | 7 | PD | none mentioned | none mentioned | varied | 100mg/hr | No side effects mentioned. |
| (Feigin et al., 2003) | 7 | PD | none mentioned | none mentioned | varied | 100mg/hr | Levodopa impaired aspects of sequence learning performance in non-demented PD patients; worsening in declarative score during motor sequence learning task suggests levodopa may have negative effects on aspects of cognitive processing linked to target retrieval. Levodopa also decreased activation of occipital association cortex during motor sequence learning. |
| (Friedhoff et al., 1963) | 11 | not given | none mentioned | none mentioned | not given | 2.5mg/kg | No side effects mentioned. |
| (Gancher et al., 1987) | 20 | PD (5 de novo, 4 stable, 11 fluctuating) | carbidopa | none mentioned | 1-4 mg/kg | 0.5-0.8 mg/kg/hr (lasting 2-5 hr) for untreated PD. For treated PD, rate approximated usual oral LD dose | No side effects reported for IV L-dopa Infusions lasting 2-5 hours. After oral levodopa, 2 of 5 de novo PD patients became nauseated (without emesis). |
| (Gancher et al., 1988) | 33 | PD (9 de novo, 7 stable responders, 17 fluctuating) | carbidopa | none mentioned | 0.8-3.0 mg/kg/hr total. (0.4 to 1.5 mg/kg/hr x 2 hr). | 1.5mg/kg/hr | No side effects mentioned. |
| (Gerstenbrand and Pateisky, 1962) | 1 | parkinsonism due to post-encephalitis lethargica | none mentioned | none mentioned | 200 mg | 100 mg/20-40 min | Increased systolic BP by 10 mmHg, mild mydriasis. |
| (Gerstenbrand and Prosenz, 1965) | 20 | PD, postencephalitic parkinsonism and vascular parkinsonism | none mentioned | Isocarboxazid (MAO inhibitor) | 50-75 mg/day for a few days, or with a few days interval between injections, up to 6-8 injections total | not given | L-dopa side effects included nausea, vomiting, blood pressure instability, and heat sensation. Subjects were pretreated with a MAO inhibitor (isocarboxazid) one tablet bid for 10-14 days. |
| (Gerstenbrand and Pateisky, 1963) | 30 | 2 with HD who had reserpine reduced parkinsonism; remaining subjects had postencephalitic parkinsonism, vascular or PD. | none mentioned | MAO inhibitors | 25-200 mg | 100-200 mg/20-30 min ("infusion"), 25-75 mg/5 min ("injection"), 100 mg cp po. | L-dopa side effects included: sensation of warmth in head, worsening of chorea in 2 HD subjects, nausea/vomiting, change in BP beyond 20 mmHg, vertigo, syncope, unpleasant sensation in head and abdomen, and urge to urinate. Subject underwent 14 days of pretreatment with MAO-inhibitors. |
| (Gillin et al., 1973) | 10 | Mild to moderate depression (4 bipolar depression, 4 unipolar affective disorder, 1 cyclothymic personality, 1 borderline personality) | carbidopa | none mentioned | 25-50 mg | 50 mg/2 min | Pre-REM infusions of L-dopa delayed the onset of REM sleep while infusion at REM onset shortened the length of the REM period. No detectable mood or side effects were noted except that three subjects had non-symptomatic reductions in blood pressure without change in pulse rate 5-25 min following the infusion. |
| (Goetz et al., 1998) | 5 | PD w/ daily visual hallucinations | carbidopa | none mentioned | 6 mg/kg (1.5 mg/kg/hr x 4hr) | 1.5 mg/kg/hr | The authors tried to intentionally produce hallucinations in patients who had daily hallucinations with their usual treatment at home. The IV doses were added to their oral medications. No patients developed hallucinations, even though baseline dyskinesias persisted during the infusions. |
| (Goldstein et al., 1999) | 6 | Healthy | none mentioned | none mentioned | 99-118.8 µg/kg (0.33 µg/min/kg x 5-6 hr) | 0.33 µg/min/kg | No side effects mentioned. Authors suggest an enzymatic gut-blood barrier for detoxifying exogenous dopamine and delimiting autocrine/paracrine effects of endogenous dopamine generated in a "third catecholamine system". |
| (Gordon et al., 2007) | 6 | healthy | carbidopa | none mentioned | infusion over 90 min (total dose estimated at ~1100mg.) | not given | No significant side effects; none of the side effects were above 1 (mild). Side effects included cold hands, mild irritability, headaches, nausea, stomach aches, but there were no significant differences between side effects reported by subjects on levodopa and those with placebo infusions. |
| (Gragnoli et al., 1977) | 25 | 8 healthy; 8 Diabetes Mellitus; 9 essential obesity | none mentioned | none mentioned | not clear, possibly 1.5 mg/kg | 1.5 mg/kg/10 min | None of the subjects suffered nausea or showed other signs of intolerance, or significant variations in blood pressure during the experiment. In diabetics and obese subjects, IV L-dopa causes a less marked HGH increase than in control subjects, with diabetics having more of an increase than obese subjects |
| (Grundig et al., 1969) | 14 | 9 PD, 5 normal | none mentioned | none mentioned | 50mg (control) to 100mg | 100 mg | No side effects mentioned. |
| (Hardie et al., 1984) | 20 | idiopathic PD | carbidopa or benserazide | Apomorphine (dopaminergic agonist) | up to 1500 mg/day | 80 mg/hr | Dystonia and chorea. 4 patients experienced significant sleep benefit. |
| (Hardie et al., 1986) | 7 | PD (on-off fluctuators) | PDI used but not specified | none mentioned | 1280 mg (up to 16 hr) | 32 - 80 mg/hr | No side effects mentioned. |
| (Hartvig et al., 1991) | 8 | Healthy | 1 subject given benserazide | none mentioned | 5.5 mg or 11 mg | 10 mg bolus | No side effects mentioned. |
| (Hashizume et al., 1987) | 6 | Healthy | none mentioned | none mentioned | 25 mg bolus | 25 mg (bolus in 20 mL saline) | No nausea (except for one patient who was given oral levodopa); authors suggest that L-dopa undergoes decarboxylation and sulfation continuously even when administered intravenously. |
| (Henry et al., 1976) | 13 | Depression, otherwise healthy | carbidopa | none mentioned | 50 mg (after a week's interval 6 pts got iv 50 mg DOPS or 100 mg L-DOPA without carbidopa) | 50 mg/5 min | No nausea, vomiting, hypertension, and "other untoward side effects". The study was designed to "avoid such peripheral side effects by pretreating the patients with carbidopa." IV levodopa was associated with reduced learning compared with chronic oral treatment and placebo infusions. No significant changes were found in heart rate/rhythm or blood pressure between levodopa and placebo. |
| (Hirano et al., 2008) | 11 | PD | carbidopa | none mentioned | not given | 0.56 mg/kg/hr | No side effects mentioned. |
| (Hirschmann and Mayer, 1964a) | 10 | PD | none mentioned | none mentioned | 25-50 mg | 50 mg | "No measurable, problematic side effects on the heart or circulation occurred with a slow i.v. injection of 25-50mg." |
| (Hirschmann and Mayer, 1964b) | 31 | 25 PD, 6 dystonia | none | MAO inhibitor | 25 - 50 mg; 25 mg/day for 21 days; proceeded to year-long weekly and then monthly injections of unspecified amount. | not stated | No side effects mentioned. |
| (Horai et al., 2002) | 1 | PD | stopped | none mentioned | 100 mg/hr x 19 days | 100 mg/hr | Total dose ≈ 45,600mg. No side effects mentioned. |
| (Ingvarsson, 1965a) | 3 | depression: long-standing, refractory (diagnosis unclear) | none mentioned | none mentioned | 10-50 mg/day for weeks | 50 mg/10 min | In one case, a sudden improvement in a concomitant asthmatic stridor was observed. "Depression" and "physical symptoms" improved in patients who were classified as depressed but may have had PD as well. |
| (Ingvarsson, 1965b) | 9 | not given | none mentioned | none mentioned | 50 mg iv | 50 mg | IV levodopa "abolishes asthmatic stridor." |
| (Jaffe et al., 1987) | 6 | PD | carbidopa | none mentioned | $\geq 2$ hours (at least 300 mg) | 2.5 mg/min | One subject had mild dyskinesia. Intravenous infusion of levodopa can affect the ERG in patients with PD, indicating that the human retina is sensitive to changes in the systemic levels of levodopa and that this drug or its metabolite cross the blood-retinal barrier. |
| (Juncos et al., 1987) | 7 | idiopathic PD | carbidopa | none mentioned | 24 hr/day x 6-13 days | ~1.5mg/kg /10 min (corrected for MW of L-Dopa instead of MW of L-Dopa methyl ester) | Motor fluctuations were markedly reduced with intravenous LDME. All patients noted an improvement in their condition during LDME treatment; reported benefits included improved sleep, attenuation of early morning akinesia or dystonia. There was no clinical or laboratory evidence of LDME toxicity. |
| (Juncos et al., 1990) | 12 | PD | carbidopa | none mentioned | 1.6 mg/kg | $7.1 \pm 7.6$ mg/hr | Dyskinesia. |
| (Ko et al., 2013) | 14 | PD |  |  | not given | not given, but see note | No side effects mentioned. Reportedly used same protocol as Mure et al. (2012) and Hirano et al. (2008) |
| (Kobari et al., 1992) | 15 | 9 PD, 6 PSP (progressive supranuclear palsy) | none mentioned | none mentioned | 1 mg/kg | 2 mg/kg/hr | No significant changes were noted in LCBF (local cerebral blood flow) after the administration of levodopa in patients with PSP. |
| (Kobari et al., 1995) | 34 | 16 idiopathic PD, 6 PSP, 5 olivopontocerebellar atrophy, 7 arteriosclerotic parkinsonism | carbidopa | none mentioned | 1 mg/kg | 2 mg/kg/hr | No significant changes in arterial blood pressure or heart rate. No side effects mentioned. Different patterns of regional CBF response to levodopa in PD vs PSP using xenon-enhanced CT. |
| (Lucas et al., 1975) | 33 | 18 healthy; 6 hypopituitarism; 9 chromophobe adenoma | none | none mentioned | 100mg | 100 mg bolus (1.5 hr after 25g arginine infusion) | No side effects mentioned. |
| (Maricle et al., 1995a) | 15 | idiopathic PD | carbidopa | none mentioned | 2 mg/kg | 1 mg/kg/hr | An elevation in mood ratings was seen for all 15 patients. (Mood ratings were an average of 40 before infusion, 60 during, and 42 after infusion). Mean anxiety decreased during the infusion (from 57 initially to 38 during infusion, and then increased to 62 after the infusion). Emotional fluctuations were seen in all patients, while only a third of the patients had a history of probable mood swings. |
| (Maricle et al., 1995b) | 8 | idiopathic PD (and fluctuating motor response) | carbidopa | none mentioned | 2 mg/kg daily x 3d | 1 mg/kg/hr | Effect on mood and anxiety was dose responsive. Six of 8 patients had mood response (increase in mood score greater than 20%) during high dose infusion. Reduction of anxiety began shortly after onset of high-dose infusion. Peak effect of anxiety occurred 30 minutes after infusion had been stopped and was followed by precipitous increase in anxiety. Patients had little insight into discrepancy between their subjective reports and how they appeared to observers during their dyskinetic and agitated, but relatively euphoric state. |
| (Maricle et al., 1998) | 18 | idiopathic PD | none | domperidone | 2mg/kg daily x 2d | 1 mg/kg/hr | No significant side effects. Authors believe, "A significant mood response after a 2-day levodopa holiday supports the hypothesis that pharmacologic tolerance may be involved in this process and that sensitization may appear after a relatively brief period of abstinence from levodopa even in the first year of levodopa therapy." |
| (Marion et al., 1986) | 3 | PD | benserazide | none mentioned | 755 - 1750 mg / 12 hr | 150mg / 10 min | No significant side effects mentioned. The patients did not experience any major discomfort or inconvenience during the course of the infusions and were pleased with their improved motor performance. Infusion were given for 6 hrs on day 1, and 12 hrs on day 2. One patient had mild dyskinesia. The number of on-off switches decreased and the duration of "on" periods increased in all three patients during the infusion periods compared to oral therapy. Intravenous infusion of levodopa (with P.D.I.) can give reproducible periods of constant mobility in selected patients for up to 5 consecutive days. One patient felt a feeling of "euphoria" after initial infusion. Another patient had a symptomatic fall of blood pressure from 140/80 mm Hg to 70/30 mm Hg when rate was at 99 mg/h of levodopa, so the infusion rate was decreased to 60 mg/h. |
| (Matussek et al., 1966) | 10 | depression and healthy subjects | none mentioned | none mentioned | 25-50 mg, 50-100 mg | not given | Headache, nausea. |
| (McGeer and Zeldowicz, 1964) | 10 | PD | none mentioned | none mentioned | not given | 5 mg/min | Three patients who were given L-Dopa intravenously experienced nausea when infusion rate increased to 5 mg/min; but all pts tolerated 2 mg/min with no noticeable side effects, except for one patient who reported light-headedness immediately following the infusion. |
| (Metman et al., 1997) | 25 | advanced PD | carbidopa | none mentioned | max dose is 45-540 mg (15-180 mg/10 min x up to 3 doses) | 180 mg/10 min | No side effects mentioned. |
| (Metman et al., 1999) | 4 | PD | carbidopa | none mentioned | 413-483 mg (64±5 mg/hr x 7 hr) | 69 mg/hr | No side effects mentioned. |
| (Metzel, 1965) | 61 | PD | none mentioned | MAO inhibitor | not given | not given | No side effects mentioned. In some cases dopa was combined with a MAO-inhibitor. |
| (Moorthy et al., 1972) | 8 | organic heart disease undergoing routine catheterization | none mentioned | none mentioned | 100-200 mg (avg 144 mg) | 200 mg/ 10 min | Nausea (5 pts), accompanied by vomiting (in 2 pts). The nausea was severe at 10-15 min after the start of the L-dopa infusion. Serious arrhythmias were not seen. Two patients had ventricular premature contractions. The effects on the cardiovascular system observed were slight. BP showed a tendency to fall in some patients during the initial 5 min after injection and to rise later to values higher than the control values. No serious complications were seen. The authors' observations seem to indicate that treatment with L-dopa is not particularly dangerous in patients with organic heart disease. |
| (Mouradian et al., 1987a) | 23 | PD | carbidopa | none mentioned | up to 11 days, 24 hr/day | 1.8 mg/kg/hr | Maximum rate provided in Juncos et al. (1990), who also give number of subjects as 28. No side effects. |
| (Mouradian et al., 1987b) | 4 | idiopathic PD | carbidopa | none mentioned | optimal dose infusion (not quantified) | optimal dose rate lasting at least 16 hours | No side effects mentioned; no cardiovascular complications. There was no discernible alteration in the motor response to intravenous levodopa at any time during the period of physical activity. |
| (Mouradian et al., 1988) | 29 | idiopathic PD | carbidopa | none mentioned | 200 mg | 200mg/ 10min | No side effects mentioned. |
| (Mouradian et al., 1990) | 12 | PD | carbidopa | none mentioned | 1.0 ± 0.1 mg/kg/hr x up to 12 days | variable; apparently up to 200mg / 10min as a loading dose | Minimal dyskinesias, with 1.0 mg/kg/hr as the dyskinesia threshold dose. |
| (Mure et al., 2012) | 8 | PD | none mentioned | none mentioned | 1.13 ± 0.41 mg/kg/hr (duration not reported) | not given | Doses titrated to achieve maximal UPDRS response without causing dyskinesia. No significant changes in regional cerebral blood flow. |
| (Nardini et al., 1970) | 17 | PD | none mentioned | none mentioned | 25 mg | 25 mg "slow infusion," 1.5 - 3 mg/kg/hr | Asthenia, insomnia, anxiety, headache, increased "tensori", restlessness, disorientation and confusion. No side effects in arterial pressure, digestive problems, liver or renal function. |
| (Nisijima et al., 1997) | 3 | Neuroleptic malignant syndrome (NMS) | none mentioned | Two patients infused with dantrolene. | 50 - 100 mg/day | not given | No side effects mentioned. Symptoms of NMS decreased dramatically. Authors write, "Levodopa, particularly in injectable form, should be more positively used for pharmacotherapy in patients with NMS." |
| (Nutt et al., 1984) | 9 | idiopathic PD | carbidopa | none mentioned | Total between 2200-7200 mg, (infusions were continued for 20-36 hours) | 110 mg/hr with carbidopa, 200 mg/hr without carbidopa | Severe dyskinesia in one patient. The patients moved around the ward and exercised freely due to intravenous L-dopa. Eating a high-protein meal during levodopa infusion is associated with a decline in the clinical response to the infused levodopa without any alteration in the plasma concentration. |
| (Nutt et al., 1985) | 9 | idiopathic PD | carbidopa | none mentioned | Max 1250 mg | 2.12 mg/kg/hr | Mild dyskinesia |
| (Nutt et al., 1988) | 8 | PD (with fluctuating response) | carbidopa | none mentioned | 0.28 - 2.54 mg/kg | 1.27 mg/kg/hr | Post-improvement worsening . Some mild dyskinesia. |
| (Nutt et al., 1992) | 27 | PD | carbidopa | none mentioned | 0, 0.4, 0.8, 1.6, 2.4 mg/kg/hr * 2 hrs | 2.4 mg/kg/hr | Mild dyskinesia |
| (Nutt et al., 1993) | 19 | PD | carbidopa | none mentioned | 33.3 mg/kg/ 21 hr | 1.6 mg/kg/h | Short infusions were well tolerated, long infusions less so. Two subjects had dyskinesia during long infusion and two others suffered from confusion, although short infusions were well tolerated by all subjects. |
| (Nutt et al., 1994) | 17 | idiopathic PD | carbidopa | none mentioned | 2 hr (average 1.96 mg/kg, max 3.2 mg/kg) | Max: 1.6 mg/kg/hr, mean: 0.98 mg/kg/hr | 2 patients developed nausea and one experienced lightheadedness (only during post-holiday levodopa infusions). In general, two-hour levodopa infusions were "well tolerated," with no medical complications during the levodopa holiday. |
| (Nutt et al., 1995) | 16 | idiopathic PD | carbidopa | none mentioned | 2 mg/kg | mean 0.98 mg/kg/hr | Some nausea and lightheadedness |
| (Nutt et al., 1997a) | 11 | idiopathic PD (and fluctuating response) | carbidopa | none mentioned | 2 mg/kg | 1.51 mg/kg/hr | Mild dyskinesia |
| (Nutt et al., 1997b) | 18 | PD | carbidopa | domperidone | 4 mg/kg total (2mg/kg daily x 2d) | 1 mg/kg/hr | Levodopa therapy was able to almost restore tapping speed to normal. |
| (Nutt et al., 2001) | 12 | idiopathic PD | carbidopa | none mentioned | 2 or 3 mg/kg | 1 or 1.5 mg/kg/hr | No side effects mentioned. Mood, anxiety, and blood pressure were measured at 30-minute intervals for 7 hours total, and there was no mention of any effects of levodopa on anxiety or blood pressure. |
| (Nutt et al., 2002) | 18 | idiopathic PD | carbidopa | domperidone | 4 mg/kg total (1 mg/kg/hr x 4 hr) | 1 mg/kg/hr | The same dose of L-Dopa produced progressively more severe dyskinesia with long-term L-dopa therapy but did not increase the duration of dyskinesia in patients. However increasing the dose of L-dopa in subjects with dyskinesia does not increase the severity of dyskinesia but does increase the duration of dyskinesia. |
| (Nutt and Nygaard, 2001) | 4 | All 4 had DRD (dopa-responsive dystonia); 2 had PD in addition to DRD | carbidopa | none mentioned | 2 mg/kg daily x 2d | 1 mg/kg/hr | No side effects mentioned. "In one subject, two doses of levodopa and a night's sleep abolished her dystonia and restored normal tapping rate." |
| (Nutt and Woodward, 1986) | 23 | idiopathic PD (and fluctuating response) | carbidopa | none mentioned | 3.0-13.2 mg/kg (0.5-2.2 mg/kg/hr x 6 hr) | 2.2 mg/kg/hr | 2 patients exhibited a brief burst of mobility and dyskinesia lasting minutes. Generally, with the onset of mobility, the patients had a brief burst of tremor, or tremor mixed with dyskinesia, and then became mildly dyskinetic. |
| (Ogawa et al., 2012) | 1 | PD | none mentioned | Dai-kenchu-tou (5-HT3 receptor agonist) | not mentioned | 75 mg/kg daily boluses, duration not reported | IV levodopa was used as a treatment for neuroleptic malignant syndrome. |
| (Oishi et al., 1996) | 20 | parkinsonism (PD, vascular parkinsonism) | none mentioned | none mentioned | 50 mg | 50 mg bolus | No side effects mentioned. |
| (Pare and Sandler, 1959) | 3 | Depression candidates for ECT who were responsive to Iproniazid | none mentioned | Iproniazid | 12.5 - 137.5 mg (25 mg - 275 mg racemic) | 275 mg bolus of DL-DOPA | No side effects mentioned. DL-DOPA was used. |
| (Pazzagli and Amaducci, 1966) | 11 | PD | none mentioned | none mentioned | 60mg, 90 mg, or 120 mg | not given | Hypotension, nausea, vomiting, somnolence, and mild sedation accompanied by feeling euphoric. |
| (Peppe et al., 1991) | 5 | PD | carbidopa | domperidone | 770 mg/day x 5 days, (given 110 mg/kg/hr x 7 hr) | 110 mg/hr (mean 70 mg/hr) | No side effects mentioned. |
| (Poewe, 1993) | not given | not given | none mentioned | none mentioned | not given | not given | No side effects mentioned. Unpublished review of IV levodopa studies with no total number of subjects given. |
| (Pullman et al., 1988) | 10 | 5 PD and 5 healthy | carbidopa | none mentioned | not given | Varied rates from high, middle, and low (actual dose not specified) | No side effects mentioned. |
| (Puritz et al., 1983) | 13 | 6 healthy; 7 progressive autonomic failure and multiple system atrophy | none mentioned | none mentioned | 99.875 mg | 1.175mg/min | No subjects experienced adverse effects during the infusion although one vomited after discontinuation of L-dopa. For one dosage and rate: change in AVP (plasma arginine vasopressin), MCP, and HR are given. No significant effects of L-Dopa on mean blood pressure in normal subjects, but lowered blood pressure of MSA patients. No effect heart rate in basal state, or on AVP levels. Author believes "L-Dopa should not be prescribed for patients with MSA." |
| (Quinn et al., 1982) | 3 | PD | benserazide | none mentioned | not given (only that treatment was given for about 8 hours at unspecified rate). | not given | No side effects mentioned. The patients with severe on-off fluctuations had dramatic benefit. Authors write, "Intravenous levodopa infusion obviously overcomes many of the problems of intermittent oral treatment." |
| (Quinn et al., 1984) | 10 | PD | carbidopa or benserazide | none mentioned | variable; highest total dose appears to be 187 mg/hr x 8.8 hr x 12 doses | 150mg bolus in $\geq 2$ subjects; all subjects received 100-200mg over 10min, then up to 187mg/hr (mean 125mg/hr) | Pulse and blood pressure fell, but to the same degree as with oral levodopa; "slight and transient" postural faintness (orthostasis); coldness of the limbs; nausea and vomiting; dyskinesias. No patient complained of palpitations during the infusions, and no arrhythmias were detected. Authors believe, "Continuous intravenous infusion of levodopa turns out to be the most effective way of abolishing the off state during a substantial period of the day." |
| (Rinne and Sonninen, 1968) | 36 | Idiopathic PD (24) and post-encephalitic PD (12) | none | none mentioned | 1.5 mg/kg | 1.5 mg/kg/10 min | Pulse and blood pressure changes were comparable between levodopa and placebo. Common adverse effects included nausea (47%), vomiting (31%), vertigo (19%), headache (33%), sweating (44%), and anxiety (22%); frequency of adverse effects not reported with placebo. |
| (Roberts et al., 1995) | 8 | normal | carbidopa | none mentioned | 50 mg | 50 mg/5 min | None mentioned. |
| (Robertson et al., 1989) | 28 | 12 healthy elderly and 16 healthy young subjects | both with and without carbidopa | none mentioned | 50 mg bolus | 50 mg bolus/5 min | None mentioned. |
| (Rodriguez et al., 1994) | 14 | Asymmetric PD | carbidopa | Domperidone. Apomorphine given subcutaneously. | 960 mg to 2200 mg (60-100 mg/hr x 16-22 hr) | 250 mg/hr (for short infusion). Up to 100 mg/h (for long infusion). | None mentioned. |
| (Rosin et al., 1979) | 1 | idiopathic PD and carcinoma of the rectum | carbidopa | none mentioned | 4320 mg highest total dose for a day (given between 1200-4320 mg/day for 7 days) | 180 mg/hr | No side effects or adverse effects: no "undue" abdominal distention, nausea, vomiting, cardiac arrhythmia, or hypotension. |
| (Ruggieri et al., 1988) | 20 | idiopathic PD | carbidopa | Domperidone | (360-1200 mg/day) x 3 days | 1200 mg/day x 3 days | The patients were given constant intravenous L-dopa infusion for 12 hours x 3days. Mild somnolence, nausea, and occasional vomiting were the only side effects reported.. There was an increase in blood pressure (probably due to domperidone). Maximum optimal drug rate ranged from 30-104 mg/hr with mean 53.5 mg/hr. |
| (Sage and Mark, 1991) | 1 | PD | carbidopa | none mentioned | 240 mg/day (during nighttime) | 30 mg/hr | No side effects mentioned. Oral carbidopa/levodopa was given during the daytime while i.v. levodopa was administered at night. Nighttime infusions produced immediate benefit of a good night's sleep, and nighttime levodopa infusions also reduced patient's daytime motor fluctuations. Authors suggest the levodopa infusion rate required to produce the best results was between 40 and 45 mg/hr. |
| (Sasahara et al., 1980b) | 5 | PD | none mentioned | none mentioned | 50 mg | 50 mg/20 min | No side effects mentioned. |
| (Schuh and Bennett, 1993) | 6 | advanced idiopathic PD | carbidopa | none mentioned | 57.6 mg/kg ,(given 24 hr/day x 3-8 days) | 2.4 mg/kg/hr | L-Dopa induced dyskinesia, but only occurs because of the progression of PD. No other side effects mentioned. |
| (Sherzai et al., 2002) | 16 | moderate to advanced PD | none mentioned | none mentioned | not given | not given | No side effects mentioned. No medically significant drug toxicity was observed. |
| (Shinoda et al., 2013) | 1 | PD | none mentioned | none mentioned | 75 mg | 50 mg bolus | Patient developed neuroleptic malignant syndrome (NMS) due to underdosing of IV levodopa as a result of dilution in extracorporeal circulation during open heart surgery. |
| (Shoulson et al., 1975) | 5 | PD | carbidopa | none mentioned | not given (duration of 3 hr at unspecified rate). | not given | No side effects mentioned. No significant changes in pulse rate or blood pressure occurred. |
| (Siddiqi et al., 2015) | 29 | Tourette syndrome and healthy controls | carbidopa | none | 0.6426 mg/kg | $2.882 \times 10^{-5} \times (140 - \text{age})$ mg/kg/min | No significant difference in pulse, blood pressure, or orthostatic change between IV levodopa and placebo when co-administered with carbidopa. |
| (Skalabrin et al., 1998) | 9 | advanced PD | carbidopa | none mentioned | not given | 2.6-3.0 mg/kg/hr | Doses escalated until a maximum of 3.0mg/kg/hr infusion rate was achieved, OR the subject experienced maximum dyskinesia, or developed nausea or hypotension. |
| (Sohn et al., 1994) | 42 | PD | carbidopa | none mentioned | 36-150 mg | 150 mg / 10 min | No side effects mentioned. |
| (Souvatzoglou et al., 1973) | 25 | healthy | none mentioned | none mentioned | 1mg, 5mg, 12.5mg, 25 mg, or 100mg | 5 mg/ml L-dopa infused, blood samples drawn at 10 min intervals over 3-4 hours. (therefore the lowest max rate possible was 5mg/ml/min) | 2 cases at 100 mg of mild nausea lasting 5-10 min. In no instance were any cardiac effects observed. Serum GH is stimulated by 25 mg IV L-dopa. |
| (Stocchi et al., 1986) | 18 | idiopathic PD | carbidopa | none mentioned | 1080–3750 mg total (360-1250 mg/day for 3 days) | 1250 mg/day x 3 days | No side-effects, except for a mild somnolence during the first day, were recorded. Blood pressure, cardiac electric morphology, and rhythm did not change significantly during the study. Authors argue intravenous infusion could be a precious form of rating the real single individual's L-dopa needs. They write, "L-dopa infusion remains a good technique in the overall evaluation of the parkinsonian patient and indispensable in particular situations like post-operative recovery and intensive care." |
| (Stocchi et al., 1992) | 9 | PD | carbidopa | none mentioned | 100 up to $\geq 600$ mg/d | 200 mg boluses or 400 mg/hr. For 3 subjects: "optimum rate" for 12 hr x 3 days | No side effects mentioned. Blood pressure and pulse were assessed every 15 minutes, and no mention was made of any changes to either blood pressure or pulse. |
| (Sunami et al., 2000) | 1 | akathisia | none mentioned | none mentioned | 25 mg/day x 8 days, followed by lower infusions | 25 mg/day | No side effects mentioned. Authors believe intravenous levodopa treatment "would be useful in reducing the persistent neurotoxicity (lethargy, hypersomnia, depression, agitation, akathisia, and confusion) associated with interferon-alpha." |
| (Takeuchi et al., 1993) | 8 | healthy | none mentioned | none mentioned | 50 mg | 50 mg for >10 min | Study of mechanisms of orthostatic hypotension in L-dopa treated PD. At rest, the systolic BP was significantly lowered by L-dopa administration, but diastolic BP, HR, and calf blood flow were not significantly altered by L-dopa administration. Spontaneous MSNA (muscle sympathetic nerve activity) was significantly higher than that before administration. Results support hypothesis that L-dopa and/or its metabolites act on peripheral blood vessels at sympathetic nerve terminal, thereby inducing orthostatic hypotension. |
| (Takubo et al., 2003) | 32 | Malignant syndrome (MS) | some subjects given unspecified PDI | none mentioned | 2440 mg/day (for patient before study began) | not given | No side effects mentioned. Suggests the following dosages of iv levodopa in the treatment of malignant syndrome: 300-600 mg/24 hr or 100-200 mg/3h three times a day. |
| (Tedroff et al., 1990) | 6 | PD and healthy | benserazide | none mentioned | 0.9 mg | 0.9 mg bolus | No side effects mentioned. |
| (Tedroff et al., 1992) | 8 | Idiopathic PD | benserazide | none mentioned | 200 mg | 200 mg/6 min | No side effects mentioned; brain uptake of [β-$^{11}\text{C}$]-L-DOPA was inversely correlated to the sum of large neutral amino acids in plasma. |
| (Tedroff et al., 1996) | 10 | PD | carbidopa | none mentioned | 3 mg/kg | 0.5 mg/kg/min bolus for 5 minutes | Before the study, one patient was excluded due to levodopa-induced nausea. Authors write, "levodopa is still the most effective symptomatic treatment for PD, and compared with the various dopamine agonists available, is well tolerated by most patients. The finding that the capacity for levodopa to produce increased synaptic dopamine levels is most profound in the more denervated regions of the striatum means that levodopa is acting preferentially at the site of dopaminergic denervation." |
| (Torstenson et al., 1997) | 10 | idiopathic PD | carbidopa | none mentioned | 5 mg/kg (2 mg/kg + 2 mg/kg/hr x 1.5 hr) | 0.5 mg/kg/min over 4 minutes as bolus; then 2 mg/kg/hr | No side effects mentioned. |
| (Tzavellas and Umbach, 1967) | 125 | PD | none mentioned | Propylhexadrine (amphetamine) | not given | not given | No side effects mentioned. Subjects received a combination of L-dopa and propyl-hexedrin (MAO inhibitor) |
| (Umbach, 1966) | not given | not given | none mentioned | Amphetamine | not given | not given | No side effects for L-dopa alone. Reported side effects are caused by combination treatment with amphetamines. L-dopa and amphetamine treatment of akinetic Parkinsonism patients with and without stereotaxic surgery. It is not clear how many were treated only with L-Dopa. |
| (Umbach and Baumann, 1964) | 35 | 30 PD, 5 controls | none mentioned | none mentioned | 100 mg in 13 patients and 100 mg in 17 patients | not given | Patients after stereotaxic surgery. Specific L-dopa side effects are not mentioned, but it is said that higher doses caused more severe side effects. |
| (Umbach and Tzavellas, 1965) | 30 | PD | none mentioned | Propylhexadrine (amphetamine) | 50 mg | not given | L-Dopa alone caused drop in BP. |
| (Verhagen Metman et al., 1998a) | 6 | idiopathic PD | carbidopa | Dextromethorphan | up to 65 ± 14mg | not given | No side effects mentioned. Brief IV infusions (10 min each, 4 hours for a total of 9-12 infusions). |
| (Verhagen Metman et al., 1998b) | 14 | PD | carbidopa | none mentioned | ≥ 150 mg | 150 mg/10 min | No side effects mentioned. |
| (Voller, 1968) | 180 | PD | none mentioned | In unspecified number of patients, MAO inhibitors (isocarboxazid 10mg TID or nialamide 25 mg BID) | 25 mg twice per week | not given | Increase of PR interval, tachycardia, sweating, nausea. All these were mild and transient so that no experiment was interrupted. |
| (Worth et al., 1988) | 6 | healthy | none mentioned | none mentioned | 840 µg/kg | 7µg/kg/min | Mean PRA fell by 50%; significant increase in urinary sodium excretion and effective renal plasma flow; mean diastolic blood pressure fell with no reflex tachycardia. Mean diastolic pressure fell on infusion of L-dopa. Trends towards fall in mean systolic pressure and rise in mean pulse rate on infusion of L-dopa, but these were not significantly different from changes occurring on saline infusion. |
| (Zsigmond et al., 2012) | 10 | PD | none | none | 281.25 mg | 375 mg/hr for 45 minutes | No side effects mentioned. In 2 patients who had previously discontinued oral levodopa/carbidopa due to nausea, high doses of IV levodopa were well-tolerated and relieved symptoms. |
| **total references 142** | **Total 2776** |  |  |  |  |  |  |

Concomitantly administered peripheral decarboxylase inhibitors included carbidopa and benserazide. Often, PDIs affected clearance and volume of distribution (as mentioned above), minimized gastrointestinal symptoms, and allowed subjects to be given lower doses of levodopa. Other cconcomitant drugs were also listed, to help explain any side effects that might be caused by concomitant drug administration, or an interaction with levodopa, rather than by levodopa alone. These included adenosine receptor antagonists (istradefylline, tozadenant [SYN115], aminophylline, caffeine), stimulants (amphetamines, methylphenidate), dopamine receptor agonists (apomorphine, terguride, SKF38393), monoamine oxidase (MAO) inhibitors, dextromethorphan, estradiol, paroxetine, and dantrolene.

A variety of neurological, psychiatric, cardiovascular, and other physiological effects of levodopa were monitored (see Table 1). There were no reported deaths. There were no instances of psychosis, even when attempting to elicit it in susceptible subjects (Goetz et al., 1998). There were also no life-threatening events (serious adverse effects) following intravenous levodopa administration at high doses, regardless of whether a PDI was co-administered. With co-administration of a PDI, the dosage range causing side effects (mainly nausea and asymptomatic hypotension) was a 0.5-2.0 mg/kg/hr infusion or a 45-150 mg bolus. Without a co-administered PDI, side effects were reported at a 1.5-3.0 mg/kg/hr infusion or a 60-200 mg bolus. It should be noted that occurrence of side effects was more likely with higher doses, but other factors such as age, sex, disease severity, and prior treatment also played a role in side effects of levodopa.

Other than these side effects found at high doses, several milder or less frequent side effects were reported. These primarily included mild autonomic changes (orthostasis and tachycardia), psychiatric changes (sedation, anxiety, insomnia, and improvement in mood), and neurologic effects (improvements in tics, REM sleep changes, subjective weakness, headaches, and increased dyskinesias). Various other effects were noted in isolated reports (listed in Table 3). It is important to note that both side effects and efficacy depended strongly on subject factors including gender, age, past treatment, and disease state. Also, dsykinesia was mentioned as a side effect only in patients with PD, and most often in those with a long history of previous levodopa treatment.

Motor benefits of levodopa in PD have been demonstrated conclusively. Additional reported benefits of intravenous levodopa treatment in PD included improved sleep (Hardie et al., 1984) and attenuation of early morning akinesia or dystonia (Juncos et al., 1987). In other patient groups, benefits of intravenous levodopa included improvement of the comatose state in hepatic encephalopathy (Abramsky and Goldschmidt, 1974) and improvement in depressive and somatoform symptoms (Ingvarsson, 1965a). One report found it more effective than dantrolene for treating neuroleptic malignant syndrome (Nisijima et al., 1997). More recently, intravenous levodopa treatment was found to alleviate the neuropsychiatric adverse effects associated with interferon-alpha (lethargy, hypersomnia, depression, agitation, akathisia, and confusion) (Sunami et al., 2000).

## Discussion

The existing literature strongly supports the safety of intravenous levodopa, which has been used in humans for more than half a century (Pare and Sandler, 1959). Intravenous levodopa has been administered to over 2600 human subjects. Despite infusion rates as high as 5.0 mg/kg/hr and boluses as large as 200 mg, there are no recorded instances of death or of other serious adverse effects of intravenous levodopa, nor have there been documented cases of other serious side effects, such as psychosis, that might limit its use in humans. Milder side effects, the most significant of which are nausea and vomiting, were most prominent with rapid infusions in the range of 1-2 mg/kg or 100-200 mg over less than 15 minutes (Bruno and Bruno, 1966; Fehling, 1966; Rinne and Sonninen, 1968; Moorthy et al., 1972; Quinn et al., 1984; Black et al., 2003).

These conclusions are supported by safety data from other species. The Registry of Toxic Effects of Chemical Substances reports the lowest published toxic dose of levodopa in any non-human species as 2.5 mg/kg, referring to a subtle behavioral effect on a learning measure in a mouse (NIOSH and Biovia, 2015).^*^ The lowest intravenous levodopa dose that was lethal to half of subjects (LD50) was “ >100 mg/kg” in rats. In mice, the LD50 ranges from 450 mg/kg (administered intravenously) to 4449 mg/kg (administered subcutaneously). Typical human doses are in the range of only 1 mg/kg; thus, human studies with intravenous levodopa administer doses substantially lower than those dangerous to nonhuman mammals.

Intravenous levodopa has similar efficacy and side effects as oral levodopa (Connolly and Lang, 2014) and dopamine agonists (Bonuccelli and Ceravolo, 2008). These include GI (nausea, vomiting, and abdominal discomfort) and neuropsychiatric effects (sedation, dyskinesias). Nausea and orthostatic hypotension, side effects of both IV and oral levodopa, are largely blocked by PDIs and are less common in patients accustomed to dopamimetic treatment. The other side effects are infrequent and neither serious nor life-threatening (Connolly and Lang, 2014). When given with adequate PDI pretreatment, intravenous levodopa has minimal if any cardiovascular effects (Siddiqi et al., in press).

The safety of IV levodopa is important for patients but also for regulatory review. Changing the route of administration of any drug in a study traditionally necessitates submitting an IND application if changing the route of administration “ significantly increases the risks … associated with the use of the drug product” (<u>§21</u> <u>CFR 312.2(b)(iii)</u>). The data from our review of the literature suggest that intravenous administration of levodopa does not significantly increase the associated risks of levodopa in comparison to oral administration. In summary, studies conducted throughout the past half century support the safety of IV levodopa administration in human patients.

## Acknowledgments

The authors gratefully acknowledge the assistance of Claire Devine, J.D. (former affiliation: School of Arts and Sciences, Washington University in St. Louis) and of Beth Beato.

Manuscript preparation was funded in part by the National Institutes of Health (K24 MH087913).

An early summary of this work was presented at the World Parkinson Congress, Washington, DC, USA, February, 2006; the poster is <u>archived at F1000 Posters 6:268, 2015</u>.

## Author contributions

Literature search: NKA, SHS, CLG, KJB

Writing: SHS, CLG, JSP, KJB

Statistics: NKA, KJB

Translation from German: MK

## Footnotes

* RTECS reported the lowest toxic dose as “ 100μg/kg,” but the dose in the cited reference was actually 100μg/g = 100mg/kg: Takahara J, Yunoki S, Hosogi H, Yakushiji W, Kageyama J, Ofuji T (1980) Concomitant increases in serum growth hormone and hypothalamic somatostatin in rats after injection of γ-aminobutyric acid, aminooxyacetic acid, or γ-hydroxybutyric acid. Endocrinology 106:343-347.

